# The impact of western versus agrarian diet consumption on gut microbiome composition and immune dysfunction in people living with HIV in rural and urban Zimbabwe

**DOI:** 10.1101/2025.07.18.665619

**Authors:** Angela Sofia Burkhart Colorado, Nichole M Nusbacher, John O’Connor, Tyson Marden, Janine Higgins, Charles Preston Neff, Suzanne Fiorillo, Thomas B Campbell, Margaret Borok, Kathryn Boyd, John Sterrett, Brent E Palmer, Catherine Lozupone

**Author notes:** corresponding author E-MAIL ADDRESSES: Catherine Lozupone. authors contributed equally (Angela Sofia Burkhart Colorado, Nichole M Nusbacher).

## Abstract

**Background:** People living with HIV (PLWH) suffer from chronic inflammation even with effective antiretroviral therapy (ART). A high-fat, low-fiber western-type diet has been linked with inflammation, in part through gut microbiome changes. In sub-Saharan Africa (SSA), a region with high HIV burden, urbanization has been linked with a shift from traditional agrarian towards westernized diets, and with changes in food security. To explore the relationship between diet, inflammation, and the gut microbiome in PLWH, we enrolled 1) ART Naïve PLWH who provided samples before and after 24 weeks of ART, 2) PLWH on ART at both timepoints and 3) HIV-seronegative controls. Individuals were evenly recruited from rural and urban Zimbabwe (locations were 145 kilometers/90 miles apart). Using a food frequency survey designed to measure intake of agrarian versus western-type food items in Zimbabwe, we determined how diet differs with urbanization, HIV-infection and treatment, and is related to inflammation and the gut microbiome.

**Results:** Individuals residing in a rural area of Zimbabwe less frequently consumed high-fat, low-fiber western type food items and had lower consumption of diverse food items overall, except for sadza-a subsistence staple-processed from home-grown grains. Consumption of a more western-type diet correlated with lower CD4+ T cell percentage in untreated and treated PLWH and with increased T cell exhaustion in PLWH on ART. PLWH on ART at time of enrollment also consumed diverse food items at a lower frequency and more often were underweight. Low food consumption correlated with muted improvements in T cell exhaustion after 24 weeks of ART. Individuals residing in the rural area had more *Prevotella*-rich/*Bacteroides*-poor microbiomes, but this did was not significantly mediated by diet. western diet consumption reduced the diversity of carbohydrate substrate degradation capabilities in the microbiome, based on predictions made using metagenomic polysaccharide utilization loci.

**Conclusions:** Taken together, this work supports that consumption of more high-fat/low-fiber type food items has the potential to exacerbate HIV pathogenesis in a sub-Saharan setting where HIV burden is high and reinforces the importance of nutritional support for promoting immunologic response to ART in PLWH in SSA.

## Introduction

Human Immunodeficiency Virus 1 (HIV-1) infection is characterized by decreased numbers of CD4+ T cells, chronic immune activation and inflammation that is only partially remediated with successful antiretroviral therapy (ART) [1]. Understanding factors related to chronic immune activation and associated co-morbidity is essential for devising strategies to protect the health of people living with HIV (PLWH). Consumption of “westernized” diets that are high in fat, refined grains and processed foods has been linked with inflammation and subsequent development of diseases including diabetes and cardiovascular disease [2]. These effects have been linked at least in part with changes in gut microbiome composition [3]. The relationship between diet and inflammation in PLWH has not been deeply explored, although studies have reported an association between consumption of western-type diets (higher in fat and/or sugar and lower in fiber) and lower CD4+ T cell count in PLWH on ART in Iran and Europe [4, 5]. Consumption of a diet high in saturated fat and cholesterol by Simian immunodeficiency virus (SIV)-infected non-human primates also promoted increased systemic immune activation, inflammation and SIV-disease progression [6, 7]. This was accompanied by an altered gut microbiome composition, gut damage, and microbial translocation [6].

About 70% of the global HIV epidemic is concentrated in sub-Saharan Africa (SSA) [8]. A traditional diet in SSA is high in fiber and low in fat and sugar and processed foods [9], which would in principle support positive health outcomes in PLWH. However, there has been a transition in middle- and low-income countries to the consumption of more “western” type foods that are high in saturated fats, refined grains, and processed foods, and low in dietary fiber [10]. This has been associated with increased prevalence of non-communicable diseases that are disproportionately present in developed countries, such as obesity and cardiovascular diseases [10]. Studies examining these nutrition transitions in SSA also show that they can influence food security and malnutrition-related disease (associated with developing societies) in complex ways [9, 11–13].

In previously published work, we characterized differences in inflammatory markers (IL-6, c-reactive protein (CRP)), and T cell exhaustion, activation, and gut homing in individuals living in urban and rural Zimbabwe [14]. As expected, we found elevated levels of inflammation and T cell activation and exhaustion with untreated HIV infection. Although expected reductions in IL-6, T cell activation, and exhaustion were observed with ART-induced viral suppression, these improvements were more pronounced in those living in urban versus the rural area [14]; and we did not interrogate the role that diet may play in this differential response to ART or in immune phenotypes or microbiome composition.

In this work, we proposed that PLWH consuming a more western-type diet may be particularly vulnerable to chronic inflammation compared to those consuming more agrarian diets, and that the gut microbiome would potentially play a mediating role in these relationships. To test this hypothesis, we evaluated the consumption of western versus agrarian foods in PLWH and seronegative controls from rural and urban areas in Zimbabwe. We then related diet to inflammatory immune activation phenotypes and gut microbiome composition.

## Results

### Participant Recruitment

A total of 168 participants were recruited from two clinic locations, one rural and one urban, with 153 completing all study visits. The rural recruitment site was the Mutoko District Hospital, a hospital located in Mutoko (population approximately 12,500) servicing surrounding rural villages and about a 2-hour drive (90 miles) from the city of Harare. The urban individuals were recruited from the Mabvuku Polyclinic, a large urban clinic administered by the City of Harare. Individuals were recruited across 3 cohorts: 1) PLWH who were not on ART at the first timepoint but who subsequently commenced first-line ART with efavirenz/lamivudine/tenofovir disoproxil fumarate (EFV/3TC/TDF) and the prophylactic antibiotic cotrimoxazole (ART Naïve), 2) PLWH who were on this same ART regimen and cotrimoxazole at both timepoints (ART Experienced), and 3) people without HIV, hereafter referred to as healthy controls (HC). Participants were excluded from all cohorts if they had used antibiotics (apart from co-trimoxazole) within the prior two months, were pregnant, had a Body Mass Index (BMI) greater than 29.9 kg/m2, or were <18 years old. Of the 168 recruited participants, 30 were subsequently excluded from specific cohorts as detailed in **Figure S1**. Other reasons for exclusion included those with unsuppressed virus following ART and ART naïve who had no viral load at the first clinic visit (suggesting questionable ART status or HIV infection).

### Study Population

Three cohorts were evenly recruited from the rural and urban clinics and each cohort had ∼50% women. The ART experienced PLWH were significantly older than the ART naïve and HC cohorts (**Table 1**). The ART experienced cohort had a significantly lower BMI than the ART naïve and HC cohorts with more individuals in the underweight categories (**Table 1**). As we reported previously [14], a gender stratified analysis revealed that this was driven by males.

**Table 1:** Clinical and demographic characteristics of the study population by cohort at the baseline visit. P-values were calculated using the Mann Whitney U test. BMI categories were determined using World Health Organization (WHO) standards [24]. NA represents values not applicable due to a particular cohort. Values are reported as the median with the minimum and maximum range in brackets.

|  | ART Naïve PLWH<br>(Naïve)<br>(N=65) | ART Experienced PLWH<br>(Exp)<br>(N=33) | Healthy Controls<br>(HC)<br>(N=40) | Naïve vs<br>Exp | Naïve vs<br>HC | Exp vs HC |
| --- | --- | --- | --- | --- | --- | --- |
| **Arm** |  |  |  | 0.835 | 0.9423 | 0.9032 |
| Rural | 33 (50.8%) | 16 (48.5%) | 20 (50.0%) |  |  |  |
| Urban | 32 (49.2%) | 17 (51.5%) | 20 (50.0%) |  |  |  |
| **Sex** |  |  |  | 0.6159 | 0.6623 | 0.9387 |
| Female | 37 (56.9%) | 17 (51.5%) | 21 (52.5%) |  |  |  |
| Male | 28 (43.1%) | 16 (48.5%) | 19 (47.5%) |  |  |  |
| **Age (years)** | 34 [19, 51] | 46 [19, 64] | 35.5 [18, 67] | <0.001 | 0.5904 | <0.001 |
| **BMI (kg/m<sup>2</sup>)** | 22.5 [15.8, 29.3] | 20.2 [14.6, 27.8] | 22.4 [16.9, 29.7] | 0.0292 | 0.3729 | 0.008 |
| **BMI Category** |  |  |  |  |  |  |
| Severe Thinness <16.0 | 1 (1.5%) | 2 (6.1%) | 0 (0%) |  |  |  |
| Moderate Thinness <17.0 | 0 (0%) | 0 (0%) | 1 (2.5%) |  |  |  |
| Mild Thinness <18.5 | 3 (4.6%) | 5 (15.2%) | 1 (2.5%) |  |  |  |
| Normal 18.5–24.9 | 51 (78.5%) | 22 (66.7%) | 28 (70.0%) |  |  |  |
| Overweight ≥25.0 | 10 (15.4%) | 4 (12.1%) | 10 (25.0%) |  |  |  |
| **CD4 T cells count (cells/μL)** | 257 [10, 824] | 412 [77, 838] | NA [NA, NA] | 0.001 | NA | NA |
| **CD4 T cells (%)** | 23 [0, 48.8] | 36.7 [0, 55.7] | 54.6 [27.8, 70.4] | 0.001 | NA | NA |
| **CD4/CD8 T cell ratio** | 0.4 [0, 1.3] | 0.7 [0, 1.5] | 1.7 [0.4, 3.3] | 0.002 | NA | NA |
| **ART exposure (years)** | 0.5 [0.5, 0.5] | 7 [1.8, 21.7] | 0 [0, 0] |  |  |  |
| **Co-trimoxazole exposure (months)** | 0 [0, 7] | 84 [0, 138] | 0 [0, 0] |  |  |  |
| **Viral Load (copies/mL)** | 26491 [36, 1159529] | 0 [0, 103] | NA [NA, NA] |  |  |  |
| **Both Visit** |  |  |  | 0.001 | 0.001 | 0.687 |
| No | 21 (32.3%) | 1 (3.0%) | 2 (5.0%) |  |  |  |
| Yes | 44 (67.7%) | 32 (97.0%) | 38 (95.0%) |  |  |  |

### Food Consumption Differences by Locations and Demographic Data

We developed a Food Frequency Questionnaire (FFQ) to evaluate the degree to which individuals typically consumed more agrarian versus western food items. Foods categories within 7 broader food types (**Table 2**), were categorized as western or agrarian with western foods containing higher fat, sodium, and added sugar and lower fiber per serving, as described in the methods. Foods were grouped into the 34 categories by binning items with similar macronutrient values per serving (**Table S1**). Intake frequencies of each food category were reported as averages over the prior 6 months to account for seasonality, as described in the methods.

**Table 2:**
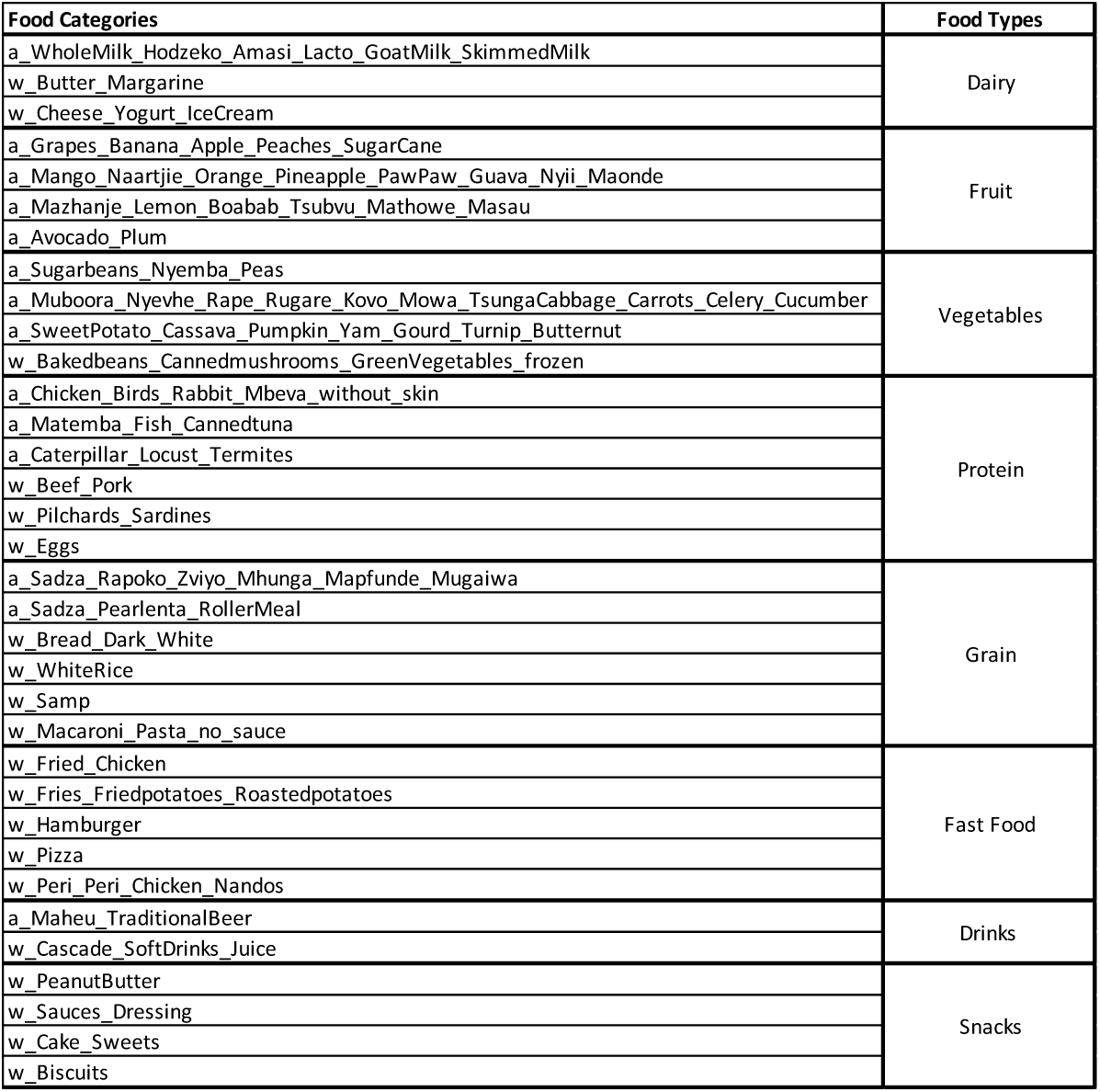
Food items surveyed in the custom FFQ. Food categories that were defined as “western” or “agrarian” types based on fat, sugar, sodium or fiber content are indicated with names beginning with “w_” or “a_” (see **Table S1**). To increase simplicity of the FFQ, foods were grouped into 34 subcategories by binning items with similar macronutrient values per serving.

To evaluate the overall trends in food consumption across individuals, we performed a Principal Component Analysis (PCA) on the frequency of the food categories consumed (**Figure 1**; **Box 1**). To facilitate interpretation of the PC axes we also plotted the broader food types in the same space using PCA (**Figure 1 C, D**). PC1 separated individuals who more frequently consumed a diverse set of foods versus those who consumed more of a local porridge called sadza, specifically one that is less refined [15]. Sadza is a filling staple made from a variety of grains (sorghum, millets, and white corn) the least refined is mugaiwa followed by roller meal then pearlenta. Rural communities tend to process (mill) their own home-grown grains yielding less refined but more nutritious sadza than commercially refined versions most widely consumed in urban areas. The FFQ surveyed participants for two types of sadza the less refined (mugaiwa) and the more refined (roller meal and pearlenta). Thus, the frequency of consumption of all broad food types were positively correlated with PC1 (hereby referred to as the Food Consumption axis; **Box 1**) as were all 34 food categories except sadza made from home-grown grains (**Figure 1C**). Food Consumption PC1 values can thus be used as an approximate estimate of the amount of diverse foods (energy) being consumed. One limitation of the study was that we were unable to get a calorie count as this would have been difficult for participants to provide, but this interpretation is consistent with Food Consumption values positively correlating with BMI at screening across individuals (**Figure S2**). PC2 and PC3 correlated to Western/Agrarian Diet spectrums 1 and 2 respectively (**Box 1**). Both spectrums differentiated individuals who frequently consumed food items that we had pre-defined as western (high values) versus agrarian (low values) (**Figure 1C** and **1D**, **Table 2**) and correlated to a Western-Agrarian Difference metric, calculated for each individual by subtracting the total frequency of agrarian foods from western foods consumed (PC2: p<2.2e^-16^, PC3: p=4.5e^-05^). Western/Agrarian Diet 1 separated individuals consuming more fast food and western proteins (high values) versus fruit and snacks (low values; **Figure 1C**, **Box 1**). Western/Agrarian Diet 2 separated individuals consuming more snack food and fast food items (high values) versus those consuming more fruit and whole grains (low values) (**Figure 1D**, **Box 1**). PC4 (hereby referred to as the Mixed axis) did not relate to agrarian versus western foods, and instead stratified individuals eating more grains and snacks (high values) and less of both western fast foods and agrarian fruits (low values) (**Figure 1D**, **Box 1**).

**Figure 1:**
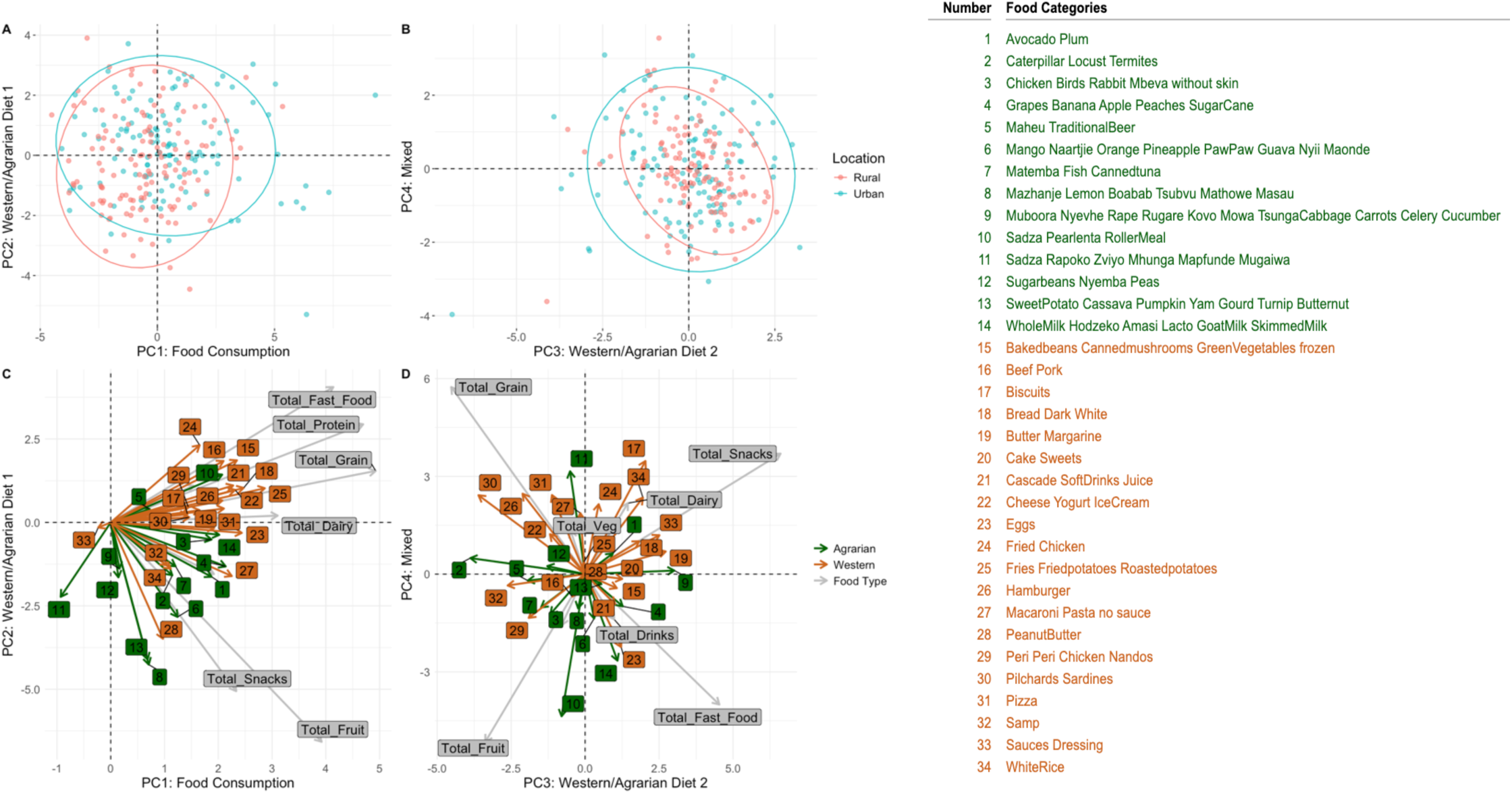
Food Category Principal Component Analysis (PCA). **(A)** Food Consumption axis (PC1) versus Western/Agrarian diet 1 axis (PC2) and **(B)** Western/Agrarian diet 2 axis (PC3) versus Mixed axis (PC4). For both **(A)** and **(B)** points and ellipses are colored according to urban or rural location. There is a 95% confidence that a point from the group (rural or urban) will fall within the region of the ellipse. Both **(C)** and **(D)** represent the same PCA spaces as A and B and arrows represent Food Categories and Food Types that correlate with the PC axes. The green and brown arrows represent Food Categories and are colored by agrarian versus western assignment of the category. Grey arrows were calculated by a PCA performed on the Food Types (Table 2). Legend for numbers is shown on right. **(C)** Food Category and Food Type PC1 versus PC2. **(D)** Food Category and Food Type PC3 versus PC4. Arrow size represents food importance, determined by Euclidean distance from the origin (larger arrows indicate greater importance).

**Box 1:** Food Category PCA Summary

| Box 1: Food Category PCA Summary |  |  |  |
| --- | --- | --- | --- |
| PC | Name | Negative | Positive |
| 1 | Food Consumption | Sadza Rich | Diverse Food Consumption |
| 2 | Western/Agrarian Diet 1 | Fruit/Snacks | Fast Food/Protein |
| 3 | Western/Agrarian Diet 2 | Grain/Fruit | Snacks/Fast Food |
| 4 | Mixed | Fast Food/Fruit | Grain/Snacks |

Participants from the rural clinic had less food consumption (lower PC1) than those from the urban clinic (**Figure 2A**). Food consumption values for the urban clinic decreased between week 0 and week 24 and increased for the rural clinic (**Figure 2A**), although there was no statistically significant difference when using linear modeling to account for changes over time, location, cohort, and water source. Individuals living in the rural location were consuming significantly more agrarian foods (p-value: 0.004) than their urban counterparts (Western/Agrarian Diet 1; **Figure 2B).** Individuals in the ART Naïve and HC cohorts had higher food consumption than the ART Experienced cohort (**Figure 2C**), which is consistent with the ART Experienced cohort also having significantly lower BMI (**Table 1**). The Western/Agrarian Diet 1 index also differed by main source of water: participants sourcing water from a bore hole or a well ate more agrarian foods as compared to those drinking from tap water (**Figure 2D**) which is consistent with individuals eating more western in urban settings where tap water is more frequently available (**Figure 2B**). Although bore holes and wells can both vary in water quality, borehole water in Zimbabwe is generally considered to be more clean and safe for drinking than community wells because of natural filtration [16]. We did not see significant differences between Western/Agrarian Diet 2 (PC3) and Mixed (PC4) PC values for cohort, location, time, and water source. Other demographic differences included a significant difference in people who worked a manual job, where those who did were more likely eat more agrarian (Western/Agrarian Diet 2) when controlling for cohort, time, and location differences (p-value: 0.029; see **Supplementary Code**).

**Figure 2:**
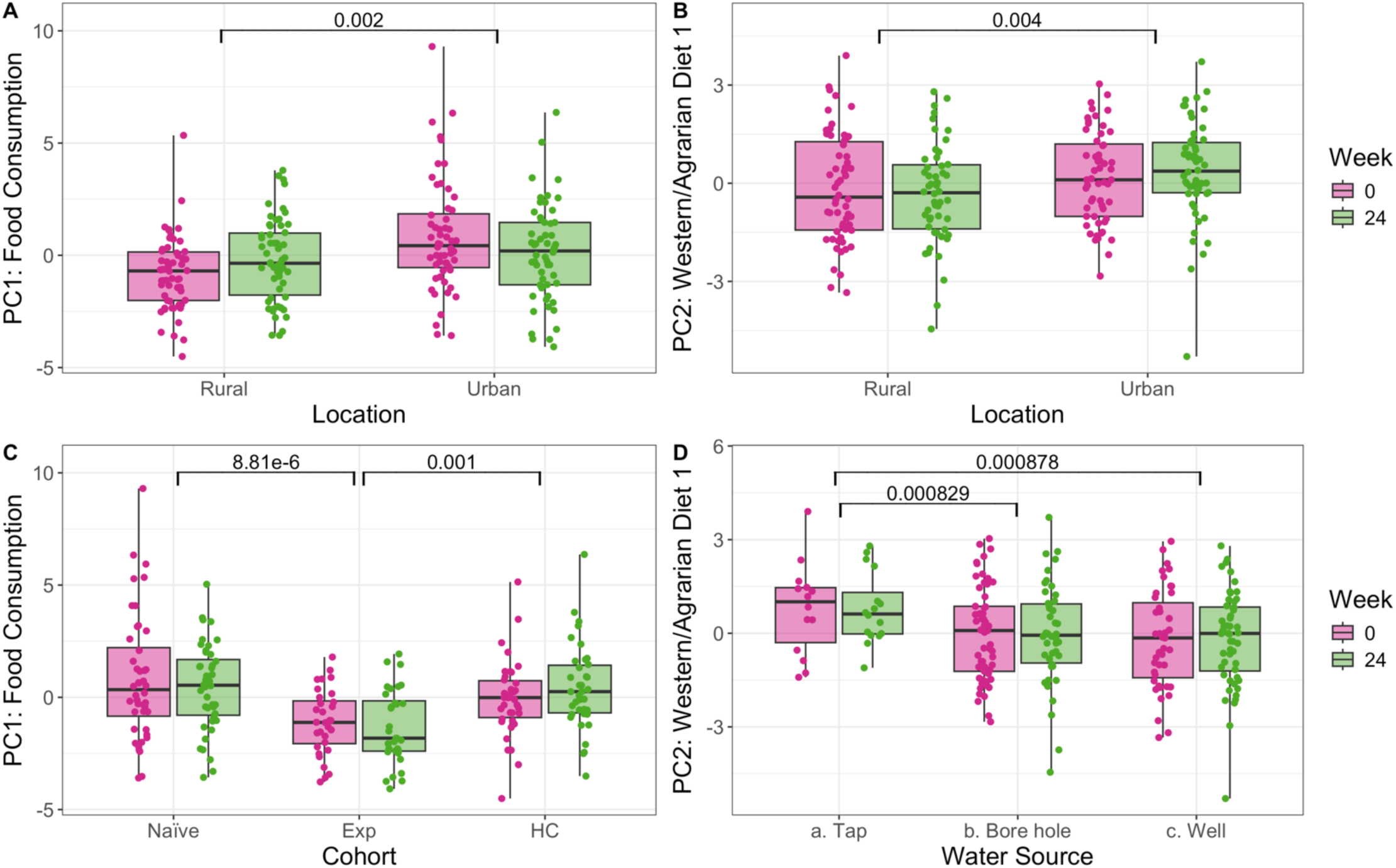
Food Category PCs Food consumption and Western/Agrarian diet 1 PCs stratified by clinic location, cohort, water source and visit. **(A)** Food consumption stratified by clinic location. **(B)** Western/Agrarian diet 1 stratified by clinic location. **(C)** Food consumption stratified by cohort. **(D)** Western/Agrarian diet 1 stratified by water source. Statistical comparisons were conducted using the following mixed linear model. Food Category PC ∼ Location + Cohort + Week + Water Source (1|PID). The model was tested for Food Category PC1, 2, 3, & 4. Only significant results are plotted here.

### Microbiome and Diet

To evaluate gut microbiome composition in this cohort, we analyzed all fecal samples with 16S rRNA targeted sequencing as described in our previous publication [14] and in the methods. We also selected a subset of 36 of the week 0 samples (ART Naïve n=16, ART Experienced n=10, Heathy Controls n=10) from the rural clinic for shotgun metagenomic sequencing that were chosen to represent individuals consuming a spectrum of diets based on Food Consumption and Western/Agrarian Diet 1 axes. Previously we used the 16S rRNA data to describe how gut microbiome composition relates to HIV-infection, ART/cotrimoxazole use, BMI and immune activation, exhaustion and inflammation across these cohorts [14]. Here, we additionally sought to evaluate the relationship between gut microbiome composition and diet. To define compositional differences in the microbiome across individuals, we performed Principal Coordinates Analysis (PCoA) on a Weighted UniFrac distance matrix [17]. The first four PCoA cumulatively explain 57.7% of the variation in the microbiome and are summarized in **Box 2 (Figure 3A** and **3B).**

**Figure 3:**
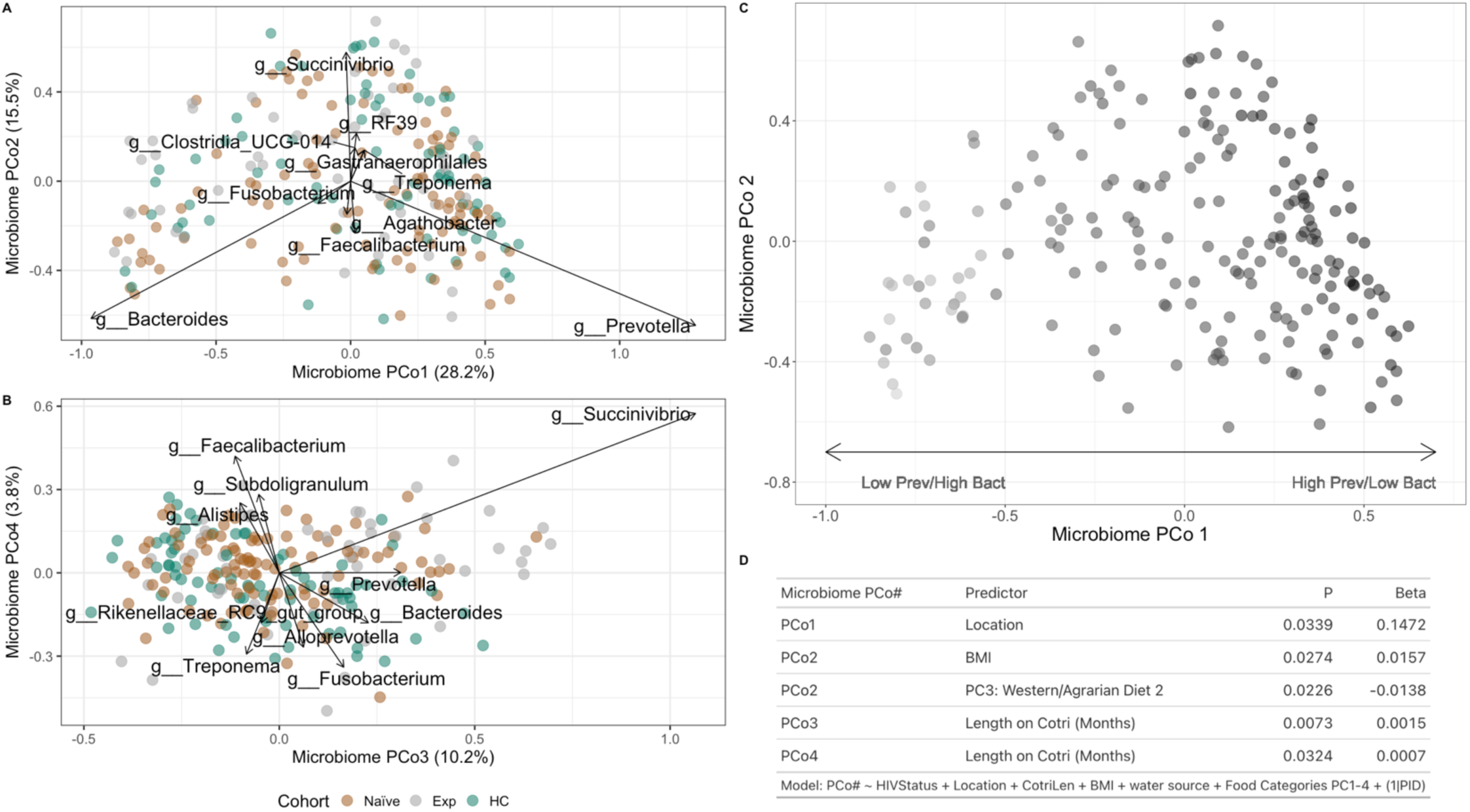
Weighted UniFrac Principal Coordinates Analysis (PCoA). **(A)** Weighted UniFrac PCoA analysis showing PCo1 versus PCo2. Points are colored by cohort. Arrow size represents genera importance, determined by Euclidean distance from the origin (larger arrows indicate greater importance). **(B)** Same as **A** except showing PCo3 versus PCo4. **(C)** Same as **A** but points are gradient colored by the log ratio of Prevotella to Bacteroides. **(D)** Results of linear modeling on Microbiome PCo 1-4. PC1-PC4. Detailed plots of significant relationships in **D** are in Figure S4.

We also sought to understand relationships between these major axes of microbiome variation (Microbiome PCo1-4, **Box 2**) and dietary intake variation patterns (Food Category PCA, **Box 1**), while considering factors that we found to influence microbiome composition in our prior publication, including HIV infection status, urban versus rural clinic location, BMI, and length of time on the antibiotic cotrimoxazole (which co-occurred with ART) using a linear mixed effects model (**Figure 3D**). We also included water source in the model since this can affect exposure to microbes. As described in our previous publication, and as shown in **Figure 3D** and **Figure S4**, factors that correlated with the first four PCoA axes of microbiome differences included whether individuals were taking the antibiotic cotrimoxazole (PCo3 and PCo4), BMI (PCo2), and residing in a rural versus urban location (PCo1). We additionally found a negative relationship between Microbiome PCo2 and the Western/Agrarian Diet 1 index, indicating some relationship between consumption of western versus agrarian food items and microbiome composition, and that a more western diet was associated communities enriched in both *Bacteroides* and *Prevotella*. Microbiome PCo 1, which strongly correlated with the *Prevotella/Bacteroides* ratio (**Figure 3C**), did not correlate with dietary patterns.

**Box 2:** Microbiome PCoA Summary

| Box 2: Microbiome PCoA Summary |  |  |
| --- | --- | --- |
| PCo | Negative | Positive |
| 1 | Bacteroides | Prevotella |
| 2 | Bacteroides/Prevotella | Succinivibrio |
| 3 | Treponema/Alistipes | Succinivibrio/Bacteroides/Prevotella |
| 4 | Fusobacterium/Treponema | Succinivibrio/Faecalibacterium |

ANCOM-BC2 [18] was used to determine differential abundance of genera with regards to cohort, Food Category PCs, and water source. There was a significant increase in genus *Fusobacterium* between the ART experienced and ART naïve cohort; and an increase in genus *Lachnoclostridium* between the ART experienced and healthy cohorts. These results are related to those of our previous study [14] but less was found, perhaps due to inclusion of different other variables in the models. While correcting for effects of cohort, no significant differences were found due to diet or water source (see **Supplemental Code**). Additionally, an Adonis test [19] was performed to assess differences in beta diversity based on diet while controlling for cohort differences, but there were none (see **Supplemental Code**).

We next used the shotgun metagenomic sequencing data that was generated on a subset of the samples to characterize the functional potential of the microbial communities. Since we were particularly interested on the effects of diet, we used dbCAN3 to annotate the capability to degrade diverse carbohydrate substrates based on the characterization of carbohydrate-active enzymes and the associated polysaccharide-utilization loci (PULs) [20]. The relative abundance of substrates across cohorts are displayed in **Figure 4A**, with the majority of carbohydrate-active enzymes targeting xylan, pectin, and host-glycan. Linear regressions revealed HIV status was associated with reduced PULs with beta-fucosidase enzymatic activity (p=0.009) (**Figure 4B and 4C**). Primary water source was associated with differences in PULs targeting beta-galactan (p=0.026) and cellulose (p=0.030) (**Figure 4B, Figure S5A**). BMI was positively associated with PULs targeting peptidoglycan (p=0.020), and length on cotrimoxazole was positively correlated with PULs acting on xylan (p=0.033) (**Figure 4B, Figure S5B**). As for dietary comparisons, all 4 dietary PC axes showed significant relationships with substrate-specific PULs, including 1) a positive relationship between Food Consumption PC1 and rhamnose; 2) positive relationships between Western/Agrarian Diet 1 PC2 and cellulose and beta glucuronan; 3) negative relationships between Western/Agrarian Diet 1 PC3 and rhamnose, sucrose, and alpha glucan; and 4) negative relationships between Mixed diet PC4 and sucrose and rhamnose (**Figure S5**). However, when looking at the breadth of microbial potential, we found that more functionally versatile microbiomes, (i.e.able to target a greater variety of carbohydrate substrates), were correlated with more fruit/grain rich-agrarian diets (lower Western/Agrarian Diet 2 PC3 index values; **Figure 4D**).

**Figure 4:**
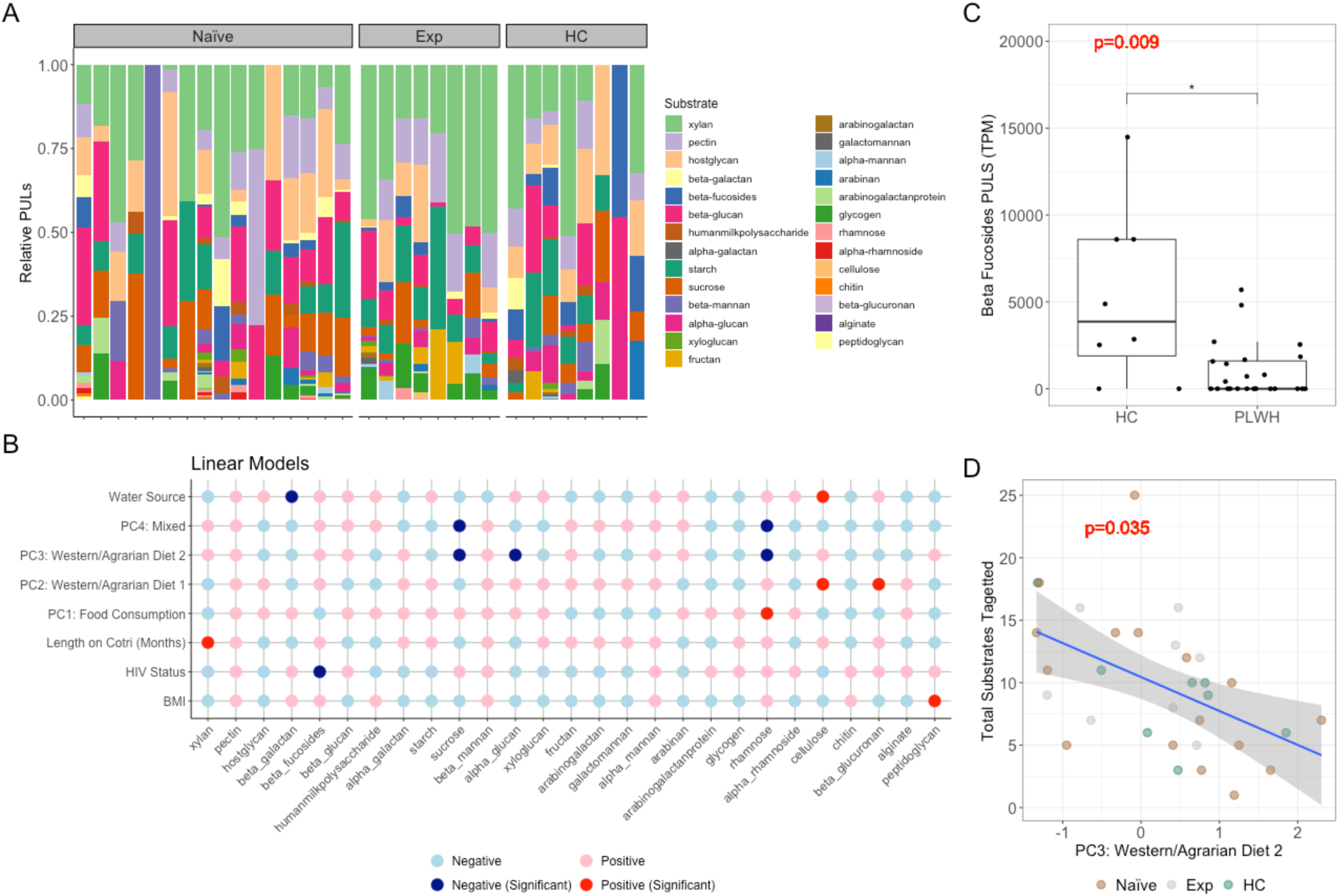
Microbial substrate targets varied by participant confounders, (**A**) Bar plot of relative polysaccharide utilization loci (PULs) in baseline fecal samples determined by shotgun metagenomic sequencing colored by substrate target and split by cohort, (**B**) Coefficients of linear models in which columns represent outcomes and rows represent predictors, predictors that had a significant impact on the model are colored red (↑/positive) or blue (↓/negative). (**C**) Boxplot of number of PULs targeting Beta-Fucosides split by HIV status. (**D**) Scatter plot of FFQ PC3 and total substrates targeted by different PULs, points colored by cohort with trend lines generated by linear regressions and p-values determined from linear models.

### Immune Markers and Diet

We next used linear regression models to test whether diet Food Category PCs were related to various immune measures. While the results of the full model are shown in **Figure S6**, we show detailed plots to highlight significant results of interest as they pertain to diet in **Figure 5**. We also included several other variables that could potentially influence immune measures in the models including age, BMI and BMI categories, education level, head of household, location, whether they performed manual chores or work, number of months on cotrimoxazole, microbiome beta diversity PCoA axes (**Figure 3**, **Box 2**), viral load, water source, and the number of years on ART. We used backwards selection [21] to define a set of predictors that associated with immune markers. Models were customized for each cohort as measurements of viral load would not apply to HC and the number of years on ART would not influence the ART-naïve cohort. Models were then applied to each timepoint and cohort separately. The relationships between immune markers and the non-diet factors are consistent with what we described in our prior publication [14] and not discussed here.

**Figure 5:**
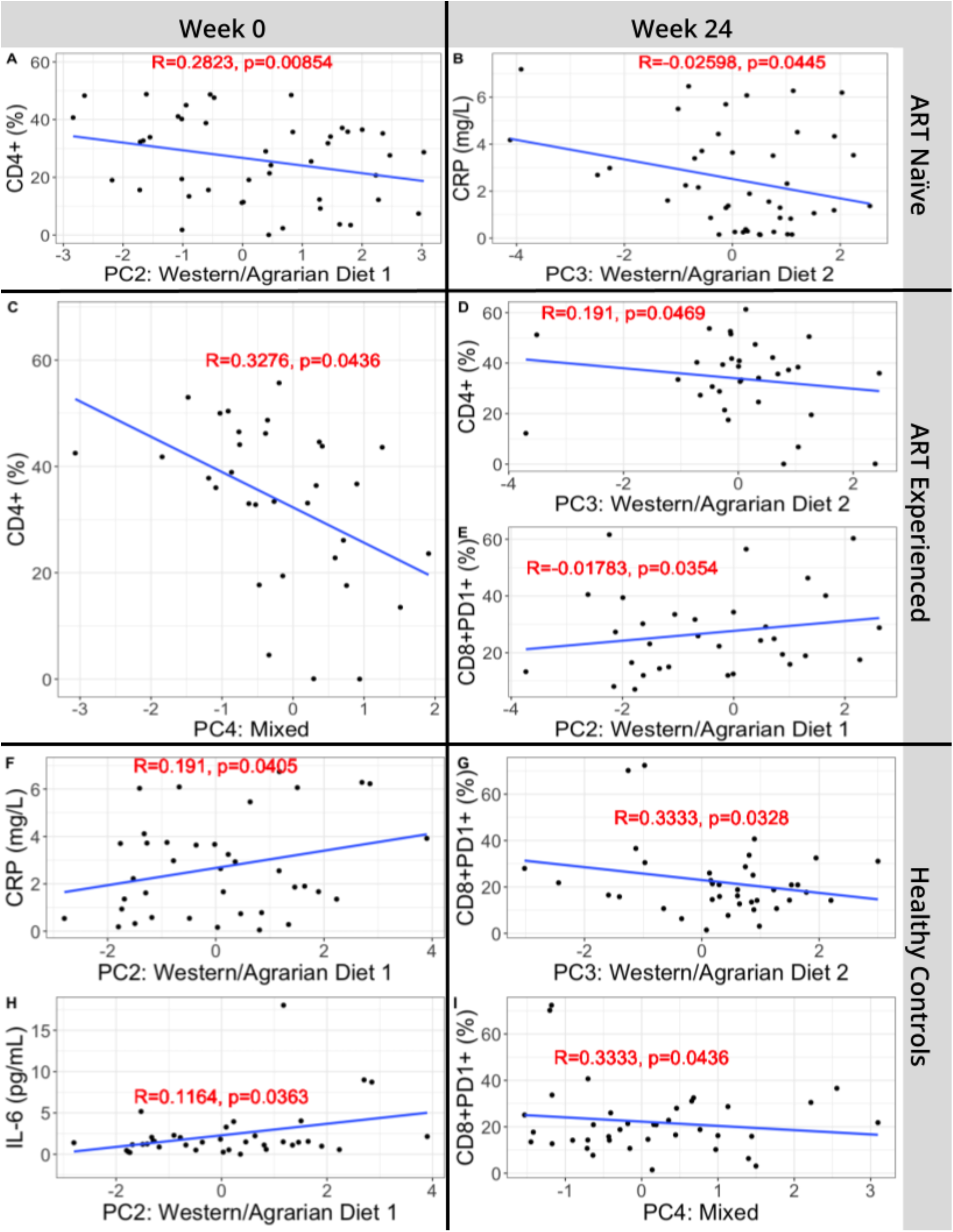
Detailed plots from Figure S5 models. R-squared and p-values in red are derived from models in Figure S6.

We observed consumption of a diet with more western fast foods/protein versus agrarian fruits (Western/Agrarian Diet 1 PC2) was associated with lower CD4+ T cell percent in ART naïve individuals (**Figure 5A**). This relationship was diminished after 24 weeks of effective ART (**Figure S6**). In the ART experienced cohort, Western/Agrarian Diet 1 PC2 correlated positively with CD8+PD1+ T cells (**Figure 5E**), showing a relationship between western diet consumption and immune exhaustion in treated infection. In the healthy controls, Western/Agrarian Diet 1 PC2 correlated positively with both IL-6 and CRP, showing a relationship between a western diet and inflammation (**Figure 5F and 5H**).

Western/Agrarian Diet 2 PC3, which is characterized by consumption of more fast food and snacks versus grain and fruit, correlated negatively with CD4+ T cell percent in the ART Experienced cohort at week 24 (**Figure 5C**), supporting a relationship between western diet consumption and low CD4+ T cell count observed previously [4, 5]. However, it was surprising that CRP correlated negatively with Western/Agrarian Diet 2 PC3 in the ART naïve cohort after 24 weeks of ART, since we expected the western diet to be associated with higher inflammation (**Figure 5B**). This could be due to patients developing what is known as immune reconstitution inflammatory syndrome, a condition observed in PLWH who have just received ART who experience tissue-destructive inflammation while they rebuild their CD4+ T cell population [22–24]. Significant differences were found when evaluating the effect of diet on CD4+CD103+ (%) and CD8+CD103+ (%) in all three cohorts (**Figure S6**), however these were primarily outlier driven. Additionally these cells are generally tissue resident cells [25], and rare in the blood which could explain why these differences appear to be outlier driven.

In our previous paper [14], we had observed that individuals in the rural clinic had a muted immunologic response to ART compared to individuals in an urban clinic. Most strikingly, individuals with untreated HIV infection at week 0 in the urban clinic had strong reductions in immune exhaustion after 24 weeks of effective ART, but individuals in the rural area did not see this expected improvement. To see whether this was related to diet, we correlated week 0 Food Category PCs with the change in CD8+PD1+ T cells with treatment. Interestingly, higher baseline values of the Food Consumption axis PC1 negatively correlated with the change in CD8+PD1+ T cells, suggesting that the lack of response to ART in the rural area could be related to nutritional status (**Figure 6**).

**Figure 6:**
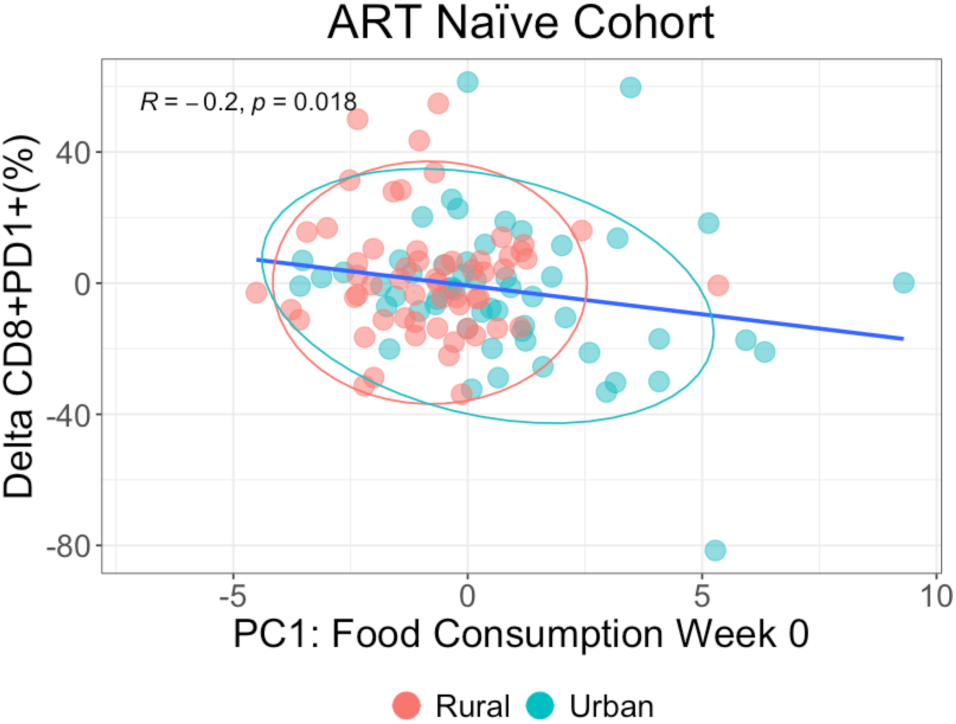
Baseline Food Category PC1 (Food Consumption) by change in CD8+PD1+ (%) over time. R and p-value calculated using a Pearson correlation. Ellipses and points colored by location. There is a 95% confidence that a point from the group (rural or urban) will fall within the region of the ellipse.

## Discussion

By integrating immune, diet, demographic and microbiome data, we were able to elucidate factors that influence the health of people living with both treated and untreated HIV infection in a rural and urban region of Zimbabwe. We developed a novel FFQ to evaluate the consumption of western versus agrarian food items in these areas. Our analysis of the FFQ data used a PCA approach to establish different dietary patterns from the data, rather than a more subjective definition of food consumption. A derived dietary pattern approach has commonly been applied [26–29], including in the context of HIV [4, 5].

We found that an important dietary pattern describing variation across this population was more frequent consumption of all types of consumed foods except for sadza made from home-grown grains, and that more frequent consumption of these diverse food items correlated positively with BMI. Since our cohort also had many underweight individuals, this food pattern was likely indicative of food security versus insecurity. Individuals from the rural area of Zimbabwe had lower food consumption compared to the urban area at the first visit (week 0), but this difference was diminished at the second visit (week 24). One potential driver of this longitudinal change was that this study was conducted during a time period in which there were steep increases in inflation rates in Zimbabwe, ranging from 3.52% to 66.8% during baseline sample collections (which were from January 2018 to March 2019) and from 4.29% to 230.54% during week 24 sample collections (which were from July 2018 to August 2019) [30], which could have particularly impacted individuals relying on store-bought foods. Financial difficulties have previously been reported to be a determinant of low fruit and vegetable intake in studies conducted in SSA [31].

We found that PLWH on ART at the time of enrollment had the lowest BMI, with 21.3% percent in the underweight categories. Given that weight loss and wasting have long been established to occur with advanced untreated HIV infection [32, 33], it is surprising that individuals in the ART experienced cohort had lower BMI at baseline than ART naive. This lower BMI was significantly associated with lower consumption of diverse foods. Lower food consumption in PLWH on ART could be due to reduced access to food while living with HIV. Indeed low diet diversity with ART in PLWH in other areas in SSA has been previously linked with low income status [34]. Another possible explanation is side-effects of ART, such as nausea or lack of appetite, but we note that these factors usually impact individuals early in their treatment course [34], and we saw no drop in food consumption in the ART naive cohort following 24 weeks of ART. In one prior study that evaluated markers of wasting to mortality in PLWH, weight loss was the strongest independent predictor of mortality, even though the analyzed population included individuals on ART [35], indicating the importance of coupling ART with nutrition support. Dietary deficiencies have also been linked with immune deficits both generally [36] and specifically in PLWH [37]. We show here that ART naïve PLWH with higher consumption of diverse foods at baseline, had larger reductions in CD8+ T cells expressing the exhaustion marker PD1 following 24 weeks of successful ART. This is likely one of the drivers of our prior description of reduced immune re-constitution with ART in the rural versus urban area [14]; poor immunologic re-constitution with ART in the rural areas may be related to individuals in those areas being less likely to have access to adequate nutrition.

Our PCA based analysis of the FFQ data also showed that the next two major patterns of diet intake variation correlated with intake of items that we had pre-defined to be “Western” versus “Agrarian” based on higher fat, sodium, and sugar and lower fiber per serving, but in different ways; PC2 differentiated people who ate more fast-food/protein versus fruit and snacks and PC3 separated individuals consuming more western-type snacks/fast-food versus fruit and grains with agrarian nutritional profiles. Individuals in the urban area were eating more fast food versus fruit compared to rural (PC2), which is consistent with other studies showing a shift towards a more western diet with urbanization in SSA [10, 38]. However, there was a strong overlap in diet in the two different areas (**Figure 1A** and **1B**).

We found that consumption of a more western diet (Western/Agrarian Diet 1 PC2) was correlated with lower CD4+ T cell percentage in untreated HIV infection. This result is consistent with a prior study conducted in SIV-infected pigtailed macaques, in which animals who consumed a diet rich in saturated fats and cholesterol experienced greater CD4+ T cell depletion during chronic untreated infection than control animals who received a regular chow diet [6]. In another study on the effects of high-fat diet (HFD)-consumption in SIV-infected rhesus macaques, a rebound in CD4+ T cell number after resolution of acute viremia was observed in control animals but not in animals fed a HFD [7]. CD4+ T cell percentage also correlated negatively with Western/Agrarian Diet 2 PC3) in individuals on effective ART. This is consistent with other studies conducted in the United States and another study conducted in Iran, that both showed sustained relationships between a western-type diet and low CD4+ T cell counts in PLWH on ART [4, 5].

Consumption of a more western diet was correlated with differences in gut microbiome composition, but not in the expected way. Specifically, since *Prevotella*-rich/*Bacteroides*-poor microbiomes have been observed in populations consuming agrarian diets and linked with fiber-rich diets in individuals in the US [39–41], we expected this microbiome signature to relate to the consumption of agrarian versus western food items, but it did not. This is the case even though individuals residing in the rural area did on average have more *Prevotella*-rich/*Bacteroides*-poor microbiomes, consistent with other studies relating this microbiome type to the degree of urbanization [38]. Since diet has previously been suggested to be a mediator of the relationship between *Prevotella*/*Bacteroides* differences across urban and rural locations, we also performed a mediation analysis to see if diet PCs mediated the relationship between location and PCo1, but no significant mediation was detected (see **Supplemental Code**). We did see some evidence of an association between microbiome composition and eating more western-type fast-food/snacks versus agrarian-type fruit/grain. Specifically, microbiome PCo2, which separated out more diverse microbiomes with higher *Succinivibrio* from less diverse microbiomes with higher *Bacteroides*, *Prevotella* and *Faecalibacterium*, correlated negatively with Western/Agrarian Diet 2 PC3 (**Figure 3**). *Succinivibrio* has also previously been reported to be increased in individuals eating more agrarian diets in Burkina Faso versus Italy [41] and in Hadza hunter-gatherers and traditional Peruvian populations [42, 43]. Microbiome PCo2 also positively correlated with BMI, suggesting that this microbiome type and associated diet can promote a healthy weight in PLWH (note that we excluded obese individuals and had a high prevalence of underweight in this study). More insight was obtained by performing shotgun metagenomic sequencing on a subset of the rural baseline samples. Notably, more functionally versatile microbiomes, which were able to target a greater variety of substrates, including rhamnose, sucrose and alpha-glucan were correlated with more fruit/grain rich-agrarian diets (lower Western/Agrarian Diet 2 PC3 index values) supporting a relationship between diverse substrate availability and functional capability of the microbiome.

## Conclusions

Taken together, this work supports that western diet consumption can advance HIV pathogenesis and promote gut microbiomes with less metabolic versatility. Consumption of less diverse food types muted immunologic improvements that are typically observed upon viral suppression with ART, supporting that nutritional support with ART is important for promoting the health of PLWH in resource-limited areas.

## Methods

### Recruitment

Study participants had a screening visit followed by two subsequent clinic visits, 24-weeks apart. During the screening visit, health/demographic information was collected including BMI, HIV infection status, and ART regimes. Clinic visits were designated Visit 1 (baseline at week 0) and Visit 2 (at 24 weeks). Trained interviewers administered the FFQ in the rural and urban locations during both study visits. Questions were also asked about hometown, education level, water sources, length of usage of the antibiotic cotrimoxazole which is used prophylactically in PLWH, occupation, transportation modes, and various daily activities.

### Fecal and Blood Specimen Collection

Fecal and blood samples were collected at both clinic visits. At the screening visit, study participants were given a fecal collection kit. Stool samples were collected in a specimen collector within 24 hours prior to Visit 1 (baseline) and Visit 2 (24 week) clinic visits, and aliquoted by the study participant into an OmniGene Gut collection system (OM-200, DNA Genotek, Ontario, Canada) for preservation of DNA. A fasting blood sample was collected by venipuncture during Visit 1 and Visit 2. Blood and fecal samples from the rural clinic were couriered to the Infectious Diseases Research Laboratory in the Internal Medicine Unit at the University of Zimbabwe Faculty of Medicine and Health Sciences (Harare), which is a 2 hour drive from the Mutoko District Hospital, in a cooler on the day of sample collection and blood was processed immediately while fecal samples were stored at -80°C. Samples for microbiome sequencing were shipped on dry ice to the University of Colorado Anschutz Medical Campus (Aurora, CO) and stored at -80 °C upon arrival.

Fecal samples were subjected to microbiome characterization with 16S rRNA (all samples) and shotgun metagenomics sequencing (subset of 32 samples). HIV positive individuals also were measured for CD4+ T cell count and viral load (**Table 1**).

### DNA Extraction and Sequencing

#### 16S rRNA Gene Sequencing

DNA was extracted using the DNeasy PowerSoil Kit protocol (Qiagen). Extracted DNA was PCR amplified with barcoded primers targeting the V4 region of 16S rRNA gene according to the Earth Microbiome Project 16S Illumina Amplicon protocol with the 515F:806R primer constructs [44]. A sterile water blank was included in each batch of extractions and PCR amplification to serve as a procedural control. Each PCR product was quantified using PicoGreen (Invitrogen), and equal amounts (ng) of DNA from each sample were pooled and cleaned using the UltraClean PCR Clean-Up Kit (MoBio). Sequences were generated on two runs on a MiSeq personal sequencer (Illumina, San Diego, CA).

#### Shotgun Sequencing

A subset of samples was sent for sequencing at the Genomics Shared Resource core at the University of Colorado. The gDNA purity, quantity and size distribution were determined with Qubit (Invitrogen) and TapeStation 4200 (Agilent) analysis prior to DNA-seq library preparation. An input of 55ng of the gDNA was mechanically sheered (Covaris) targeting 300 - 400bp DNA products and the Ovation Ultralow System V2 kit (Tecan) was used to generate DNA-Seq libraries. Paired-end sequencing reads of 150bp were generated on NovaSeq X plus (Illumina) sequencer at a target depth of 600 million paired-end reads per sample. Raw sequencing reads were de-multiplexed using bcl2fastq.

#### Blood Sample Processing

A subset of the blood sample was analyzed by flow cytometry at the Infectious Diseases Research Laboratory (IDRL) located in the Internal Medicine Unit at the University of Zimbabwe Faculty of Medicine and Health Sciences. Whole blood was collected in BD Vacutainer tubes containing sodium heparin and red blood cells (RBCs) from 500 mL of blood were lysed with 1x RBC lysis Buffer (Thermo Fisher). Cells were washed twice with staining buffer containing PBS, 2% BSA, 1mM EDTA and surface stained with BV785-labelled anti-CD3 antibody (BioLegend Cat# 317330), PerCP/Cy5.5-labelled anti-CD4 (BioLegend Cat# 317428), BV421-labelled anti-CD8 antibody (BioLegend Cat# 344748), BV605 labelled anti-CD38 antibody (BioLegend Cat# 303533), Pe-labelled anti-CD103 antibody (BioLegend Cat# 350206), Pe-Cy7 labelled anti-HLA-DR antibody (BioLegend Cat# 307616) and FITC-labelled anti-PD-1 antibody (BioLegend Cat# 329904) or appropriate fluorescence minus one (FMO) controls. Cells were washed twice with staining buffer and fixed in 1% formaldehyde. Cells were acquired on a BD LSRFortessa Flow Cytometer and analyzed by FlowJo.

Plasma was isolated and frozen for shipment to the University of Colorado for measurement of CRP and IL-6 with ELISA following the manufacturer’s protocol (CRP: R&D Systems cat. DCRP00, IL-6 Invitrogen cat. 88-7066).

Blood samples were also used to evaluate absolute CD4+ T cell count or CD4+ T cell percent using the Sysmex (formally Partec) CyFLOW Cytometer, and CD4 easy count kit or CD4 percent easy count kit following manufacturer’s instructions (Sysmex, cat: 058401, 058505, respectively), and HIV viral load using a Roche COBAS AmpliPrep /COBAS TaqMan (CAP/CTM) instrument with COBAS AmpliPrep/TaqMan HIV-1 test v2.0 kit following manufacturer’s instructions at the Infectious Diseases Research Laboratory (IDRL) located in the Internal Medicine Unit, UZ Faculty of Medicine and Health Sciences (UZFMHS).

#### Software

Unless otherwise specified, all analyses were run in R version 4.4.2 (2024-10-31) [45].

#### Data Preprocessing

Before analyses were performed, 168 participants were recruited. Of those 138 met study criteria and of those 114 participants came for both visits (**Supplemental Figure 1**). Length on ART was calculated for the ART-experienced PLWH as the number of days between the visit date and the date that they started ART. For all longitudinal analyses only study participants who provided samples at both baseline and week 24 were included.

#### Diet and Demographic Analyses

Development of the Food Frequency Questionnaire (FFQ)

Foods in the FFQ were taken from a Zimbabwean food composition database of frequently consumed foods [46], categories of consumption frequency, and the PURE/Zimbabwe Adult Semi-Quantitative FFQ [47]. Foods were grouped into: dairy, vegetables, proteins, grains, beverages, snacks, and fast food (**Table 2**). To determine if foods within each of the 7 broad food categories were agrarian or western, criteria were set according to the fat, sugar, sodium, and fiber content within each category (**Table S1**). To increase simplicity of the FFQ, foods were grouped into 34 subcategories by binning items with similar macronutrient values per serving. For example, grapes, bananas, apples, peaches, and sugar cane all had similar amounts of sugar per serving, so one frequency questions for all 5 foods had merged responses (**Table 2**). Nutritional content of individual food items were determined by using equivalent food items within the USDA Food Database (**Table S2**) [48]. Intake frequencies of each food category were scored from 0 to 6 as: Never or less than once/month, 1-3/month, 1/week, 2-4/week, 4-6/week, 1/day, greater than once/day, respectively. To account for seasonality, individuals were instructed to estimate average consumption over the past 6 months.

#### FFQ Scoring

Western-Agrarian Score: To determine if participants were consuming a more western or agrarian diet, we calculated a western minus agrarian score by adding the counts of the western foods and subtracting the total counts of the agrarian food. The more positive a W-A value, the more western foods were consumed according to the FFQ. This metric ranged from -25 to 33 with a median score of 0 for all participants: participants above 0 are considered western-dominant while participants below 0 are agrarian-dominant.

#### Statistical Analyses

Principal Component Analyses (PCAs): Two PCAs were performed on the FFQ data tables using R *stats* package [45]. The first PCA was performed on all 34 food categories (**Table 2**) and is subsequently what is used in all following statistical models. The second PCA was performed on the 8 food types (**Table 2**). The food type PCA is only used as a visual aid to help summarize the food category PCA in **Figures 1C** and **1D**.

Mixed Effects Linear Models: To evaluate the effect of location, cohort, visit and water source on the Food Category PCs the *lmer* function in the *lme4* R package [49]. Model used was:

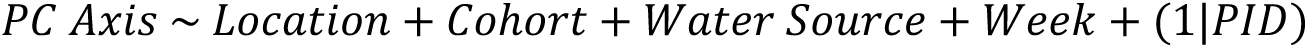

As patients came for both visits the random effect term would account for the fact that those samples would not be independent of each other. This was performed for all four food category PC axes.

#### Microbiome and Diet Analyses

Core metrics analysis and taxonomic classification

Demultiplexing of 16S rDNA gene sequences and quality control using DADA2 [50] to define ASVs were performed in QIIME2 (version 2023.5) [51]. Silva taxonomy (version 138) [52] was used to perform taxonomic classification of each ASV. Taxa that were not classified at the phylum level or that were classified as mitochondria and chloroplasts were excluded. SEPP [53] was used to produce a phylogeny of ASVs for use in diversity analyses. The feature table of ASVs was rarified to a sampling depth of 16,645 sequences per sample prior to downstream analyses.

Weighted UniFrac PCoA results from the core metrics analysis were used to calculate coordinates for biplots in QIIME2 and then visualized in R (**Figure 3**). To evaluate the effect of potential predictors on differences in beta diversity the following mixed effects models using the *lme4* package [49] were run:

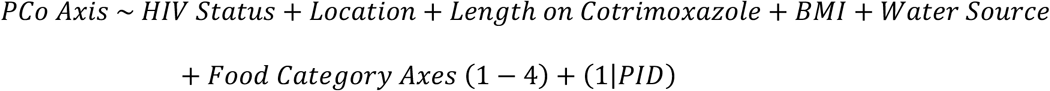

The results from these models are found in **Figure 3D** and **Figure S4**.

Substrate analysis using shotgun metagenomes

Raw metagonomic sequencing reads were trimmed with Trimmomatic under default parameters and quality was checked using FastQC and MultiQC [54, 55]. Hostile was used to filter host reads using the masked human-t2t-hla-argos985 as a human reference genome [56]. Contig assembly was performed using Megahit [57] and gene annotation was performed using Prokka [58]. Carbohydrate active enzymes, PULs, and predicted substrate targets were quantified using dbCAN3 and SAMtools [20, 59].

### Integrated Analysis of Immune Markers, Microbiome, and Diet

#### Predictive models for immune markers

To understand the impact that other variables might have on patient immunity (**Figure 5 and S6**), we curated a list of potential clinical and demographic features: cohort, location, gender, BMI, BMI categories (severe thinness, moderate thinness, mild thinness, normal, overweight, and obese), HIV status (negative/positive), week, age, education level (primary, secondar, and tertiary), water source (tap, bore hole, and well), cotrimoxazole (Y/N), length on cotrimoxazole (months), manual work, manual chores at home (Y/N), head of household (Y/N), and length on ART (years); and measures of microbial community diversity (PCo1, PCo2, PCo3, PCo4 from the weighted UniFrac PCoA). Since linear models inherently penalize large numbers of explanatory variables [60] because more variables in the model can reduce results accuracy due to overfitting, we reduced the number of explanatory variables when developing the models using backwards stepwise regression feature selection [21]. Overall, 228 samples were evaluated—including individuals who had come in at both timepoints and who had controlled infection—against the 22 aforementioned explanatory variables for each immune marker. Afterwards, viral load was included in the models pertaining to PLWH (naïve and experienced); and HIV diagnosis date was included in models pertaining only to ART-experienced PLWH. Models were created for each immune marker, cohort, and time separately using the *lm* function from stats package [45].

## Ethics and consent

### Ethics approval

The study was approved by the Colorado Multiple Institutional Review Board (COMIRB; 16-1441) guided by the principles of the Belmont Report in accordance with the US Code of Federal Regulations (CFR) 46.116, Joint Research Ethics Committee for University of Zimbabwe Faculty of Medicine and Health Sciences (UZFMHS), Parirenyatwa Hospital (JREC), Medical Research Council of Zimbabwe (MRCZ), and the Research Council of Zimbabwe.

### Consent to participate

Informed consent was obtained from all study participants.

## Consent for publication

Not applicable.

## Availability of data and material

The 16S rRNA data is available at EBI (project PRJEB66206, accession ERP151280). The shotgun sequences can be found at EBI (insert project and accession numbers, in progress). Supplemental code and corresponding data can be found on Zenodo at: https://zenodo.org/records/15843879?preview=1&token=eyJhbGciOiJIUzUxMiJ9.eyJpZCI6IjJiZDI1NDcwLTUyYzgtNDVmMC05YmJhLWYwNTBlODI5Njk2OCIsImRhdGEiOnt9LCJyYW5kb20iOiI0ZWQ3OGNlODM1MDZjOTE3ZDJiZTc3N2U2ZDAyNjNmMyJ9.clVlyx9Dh-RPD9_HrraiZNMH6lvw3LkmtMLs-2rjcxMJvqMSmHMc487-LdwLTNHZ0xKHTB3Rm_VarOXKu-kD-A.

## Acknowledgements

We would like to thank our study participants for their time and dedication to the study. We would also like to thank Mercia Mutimuri and Patricia Gambiza for support in recruitment and sample collection from study participants, Francis Jaji for study coordination support, and the staff of the Mutoko District Hospital and Mabvuku Polyclinic.

## Authors’ contributions

Angela Sofia Burkhart Colorado and Nichole Nusbacher contributed equally to this work. A.S.B.C. worked with N.N. on primary analyses, performed integrative model analyses, interpreted results, and contributed to writing of the manuscript; N.N. cleaned and processed subject metadata, managed the generation of microbiome data, and contributed to data analysis, interpretation, and writing of the manuscript; J.O’C. performed shotgun analyses and contributed to writing of the manuscript; T.M. and J.H. helped design the study, generated and interpreted the food frequency questionnaire and nutritional content of represented food, and helped N.N. develop food categories. C.P.N helped design the study, initiate the study in Zimbabwe, coordinated the receipt of primary data and samples from Zimbabwe, and generated and interpreted immune data; K.B. generated flow cytometry data in Zimbabwe and worked with C.P.N. in coordinating immune data analyses; S.F. initiated the study in Zimbabwe and interfaced directly with the Zimbabwe and US study teams to facilitate ethics approvals, recruitment, study design, and metadata storage and curation; T.B.C contributed to design and initiation of the study, interpretation of results, and editing of this manuscript; M.B. contributed to design and initiation of the study, supervision of participant recruitment and follow-up, clinical data collection, and ethics approvals; J.S. helped J.O’C. with cleaning, processing, and analysis of shotgun sequencing data. B.E.P contributed to design and initiation of the study and oversaw generation and analyses of immune data; C.L. contributed to design and initiation of the study and oversaw generation and analyses of microbiome data and integrative data analyses.

