## Supplemental Figures and Tables for "The impact of western versus agrarian diet consumption on gut microbiome composition and immune dysfunction in people living with HIV in rural and urban Zimbabwe"

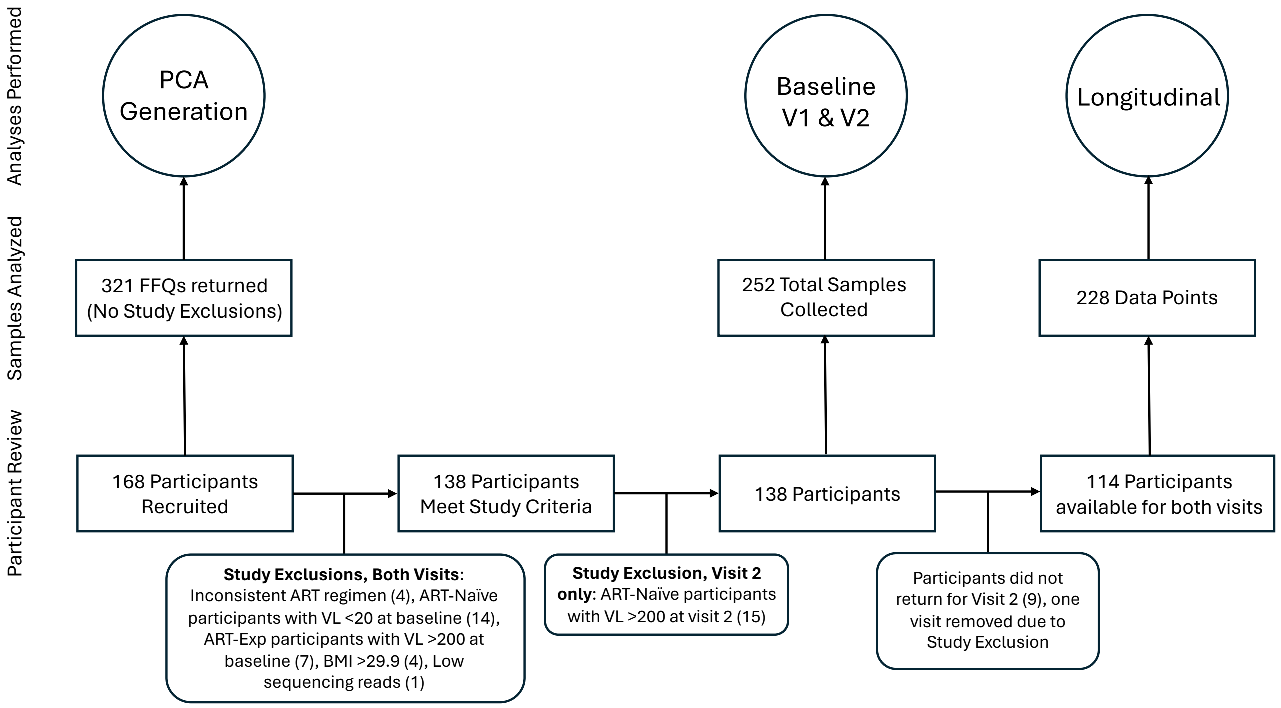


**Figure S1:** Number of study participants and samples used in different analyses and reasons for exclusion.

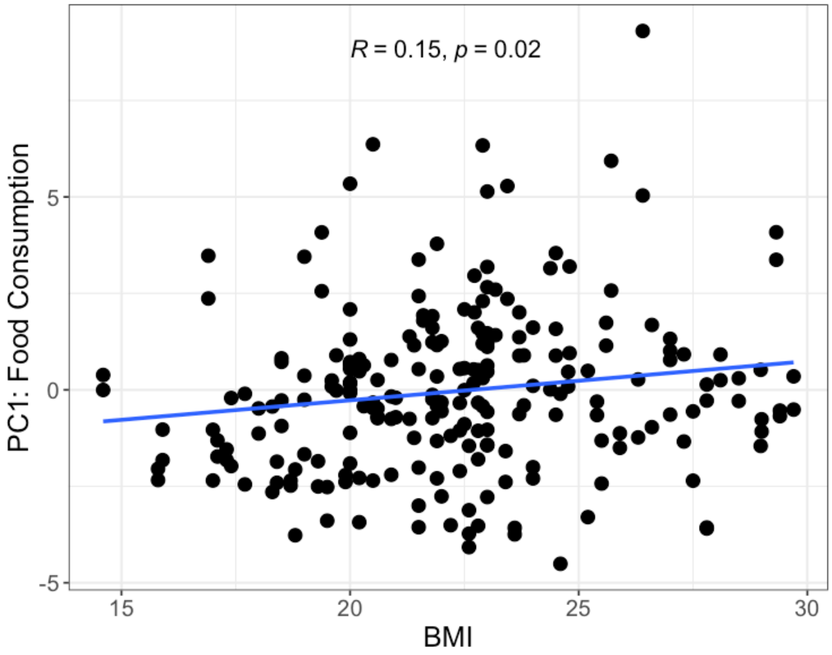


**Figure S2:** BMI by Food Category PC1 representing Food Consumption. R and p-value calculated using a Pearson correlation [58].

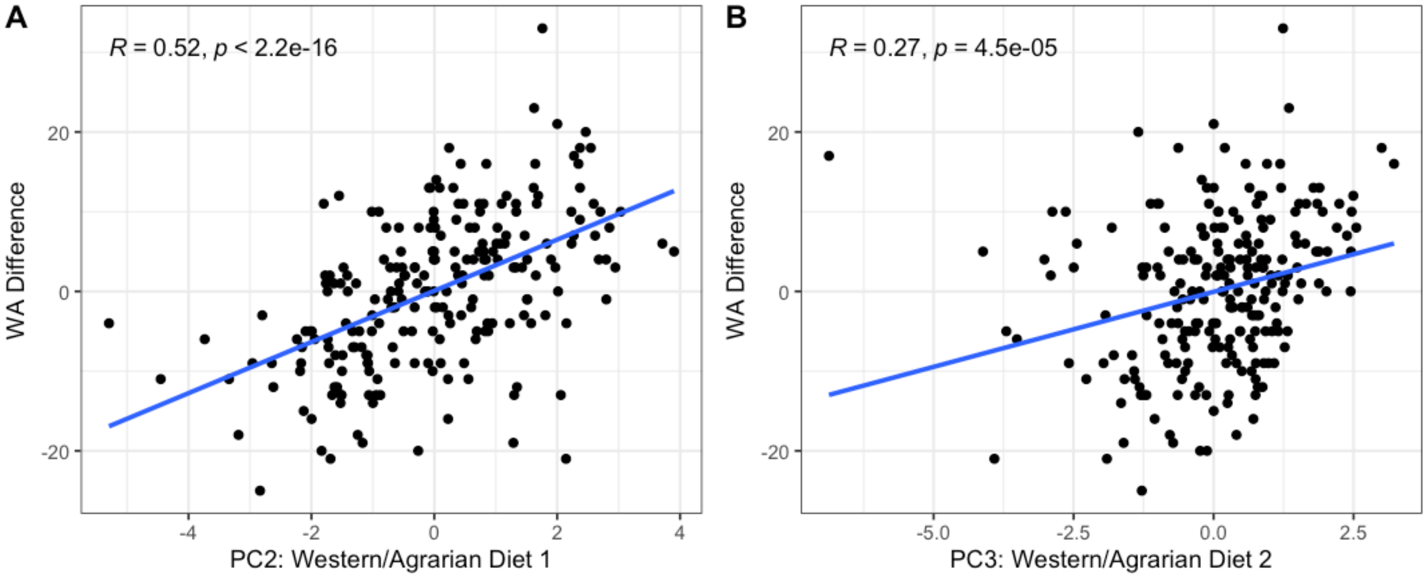


**Figure S3: (A)** correlation between Western/Agrarian Diet 1 axis from Food Category PCA with the Western/Agrarian difference score (see Methods). **(B)** Same as **A** but with Western/Agrarian Diet 2 axis. Both R and p-values are calculated using a Pearson correlation [58].

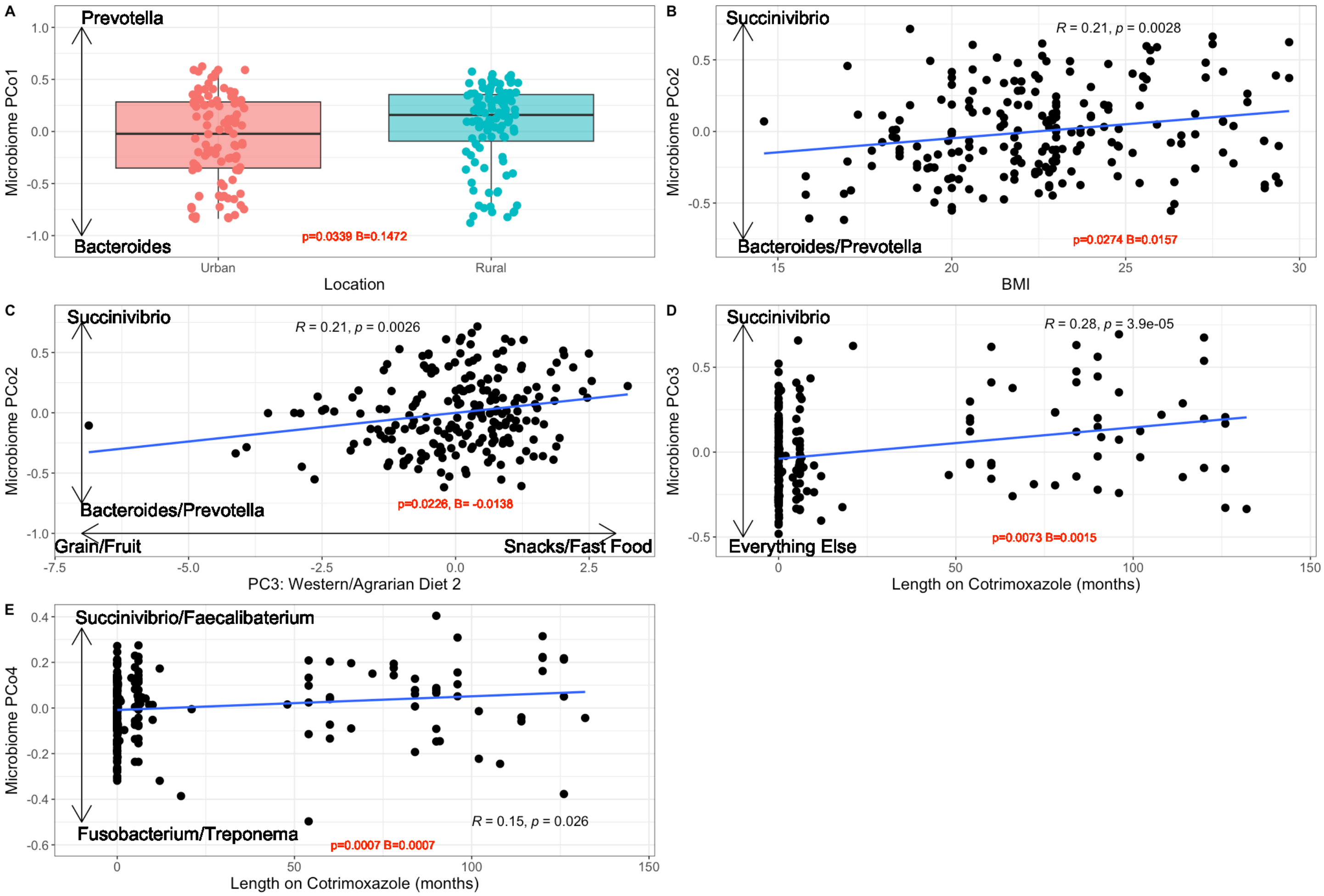


**Figure S4:** Detailed plots from Figure 3D models, p-value and betas taken from model results. P-values and R values in black derived from Pearson correlations **(A)** Microbiome PCo1 differences by location. **(B)** BMI by Microbiome PCo2. **(C)** Food Category PC3 (Western/Agrarian diet 2) by Microbiome PCo2. **(D)** Length of time on antibiotic cotrimoxazole in moths by Microbiome PCo3. **(E)** Length of time on antibiotic cotrimoxazole in moths by Microbiome PCo4.

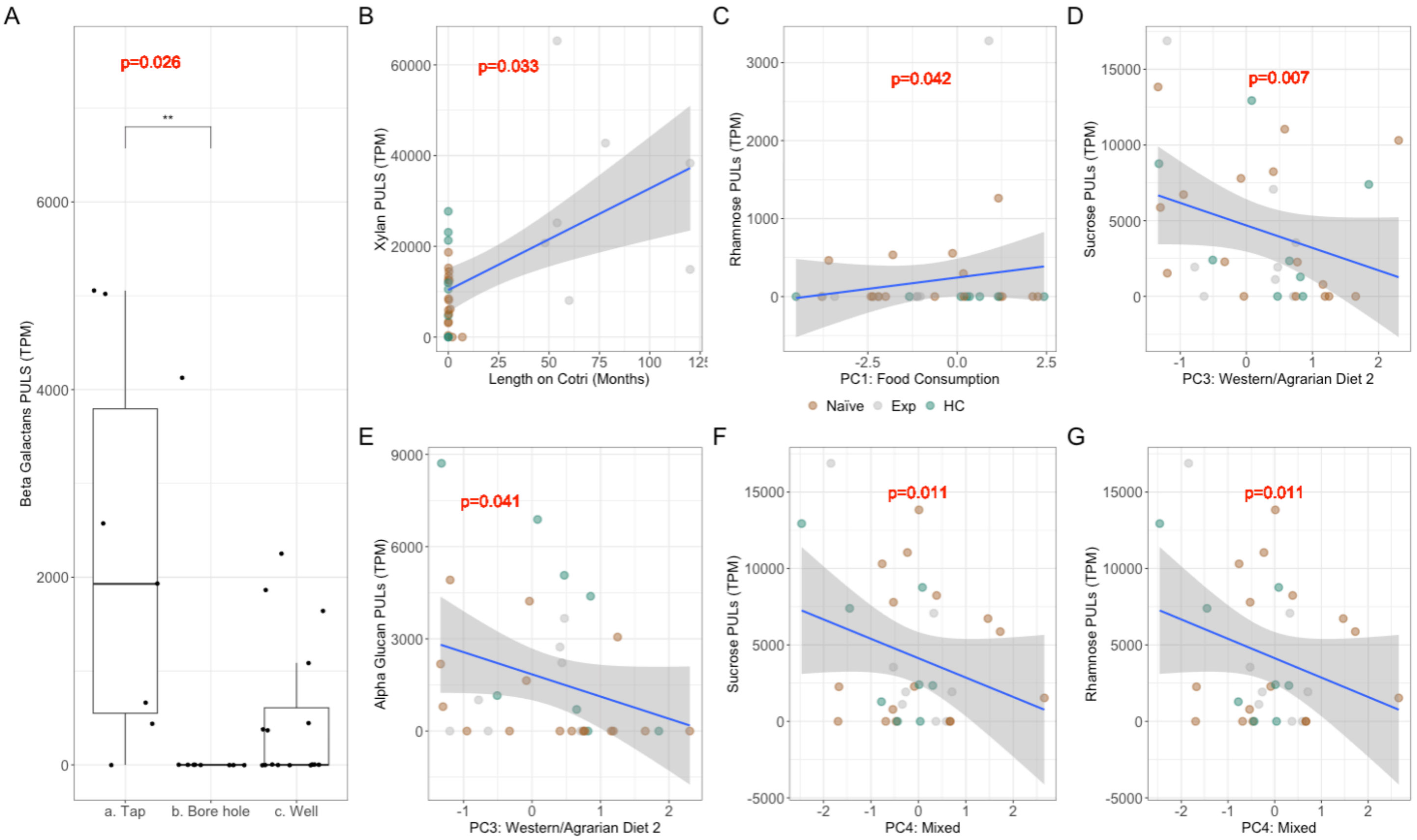


**Figure S5:** Water Source and Diet related to Utilization of Polysaccharides; (**A**) Box plot of total PULs targeting Beta Galactans split by Water Source with brackets indicating significance determined by Kruskal Wallace and Dunn's post-hoc test. **(B-G)** Scatter plots of length on Cotri and total PULs targeting xylan (**B**) FFQ PC1 and total PULs targeting rhamnose (**C**); FFQ PC3 and total PULs targeting sucrose (**D**) and alpha glucan (**E**); FFQ PC4 and total PULs targeting sucrose (**F**) and rhamnose (**G**), points colored by cohort with trend lines generated by linear regressions and p-values determined from linear models.

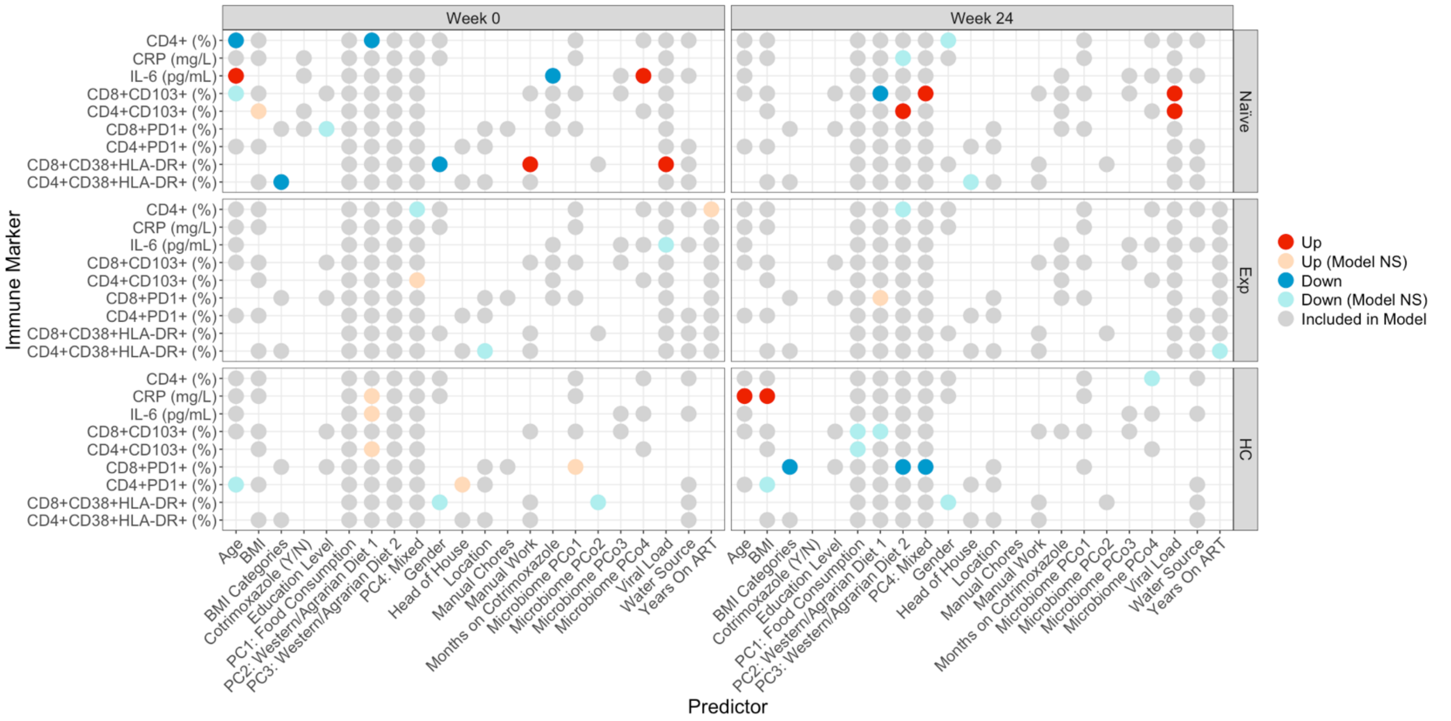


**Figure S6:** Predictive models for immune markers. Each row within each square represents one model. Circles represent predictors included in models. Predictors were determined by backwards stepwise regression feature selection. Food Category PC1-4 were retained in all models, viral load in the 2 PLWH cohorts, and Years on ART in the ART-experienced cohort. Predictors that had a significant impact on the model are colored red (↑/positive) or blue (↓/negative). Gender (up) represents higher values in males; location (up) represents higher values in the rural location; BMI categories (up) represent higher values with higher BMI; water source (up) represents higher values in people who drink from wells as opposed to tap; education level (up) represents higher values in people who went to secondary school as compared to tertiary; manual job (up) represents higher values in those who do work manual jobs as compared to those who do not. “Model NS” means that the overall model was not significant despite the predictor being significant. Gray circles represent non-significant predictors.

### Supplemental Tables

| Food Type | Fat | Sugar | Sodium | Fiber |
|---|---|---|---|---|
| **Dairy** | < 3g | < 8g | | |
| **Fruits** | | | | > 1.2g |
| **Vegetables** | | | < 250mg | |
| **Grains** | | | | > 250mg |
| **Protein** | < 3g | | | |
| **Snacks** | < 3g | < 8g | < 250mg | |
| **Fast Food** | < 3g | < 8g | < 250mg | |
| **Beverages** | | < 8g | | |

**Table S1.** Cut-off values for nutrients per serving to categorize foods as Agrarian. Any food item above or below the cutoff value for each category is categorized as western.

| Food Type | Zimbabwe Food Item | USDA Food Item |
| --- | --- | --- |
| Dairy | Margarine | Butter-margarine blend, stick, unsalted |
| | Cheese Low fat | Cheese, cottage, creamed, large or small curd |
| | Cheese Regular | Cheese, cheddar |
| | Whole Milk | Milk, producer, fluid, 3.7% milkfat |
| | Butter | Butter, whipped, with salt |
| | Yogurt | Yogurt, plain, low fat, 12 grams protein per 8 ounce |
| | Ice cream | Ice creams, vanilla, light |
| | Goat milk | Milk, goat |
| | lacto | Milk, buttermilk, fluid, cultured, lowfat |
| | Skimmed milk | Milk, fluid, nonfat, calcium fortified (fat free or skim) |
| Fruits | Grape | Grapes, american type (slip skin), raw |
| | Mango | Mangos, raw |
| | Banana | Bananas, raw |
| | Naartjie | Tangerines, (mandarin oranges), raw |
| | Apple | Apples, raw, with skin |
| | Peaches | Peaches, canned, juice pack, solids and liquids |
| | Orange | Oranges, raw, all commercial varieties |
| | Pineapple | Pineapple, raw, all varieties |
| | Paw Paw | Papayas, raw |
| | Sugar Cane | Sugar cane |
| | Guava | Guavas, common, raw |
| | Mazhanje | Loquat |
| | Lemon | Lemons, raw, with peel |
| | Baobab | Quinces, raw |
| | Avocado | Avocados, raw, all commercial varieties |
| | Plum | Plums, dried (prunes), stewed, with added sugar |
| | Maonde | Fig |
| | Nyii | Date |
| Vegetables | Beans | Beans, shellie, canned, solids and liquids |

| | | |
|---|---|---|
| | Mushroom, canned | Mushrooms, canned, drained solids |
| | Mushamba | Pumpkin, canned, with salt |
| | Green vegetables | Vegetables, mixed, frozen, cooked, boiled, drained, with salt |
| | Beans | Beans, snap, green, canned, regular pack, drained solids |
| | Carrot, fresh | Carrots, raw |
| | Turnip | Turnips, cooked, boiled, drained, without salt |
| | Cabbage, fresh | Cabbage, raw |
| | Lima Beans | Lima beans, immature seeds, canned, no salt added, solids and liquids |
| | Peas | Peas, green, raw |
| | Celery | Celery, raw |
| | Okra | Okra, frozen, cooked, boiled, drained, without salt |
| | Mushrooms, fresh | Mushrooms, white, raw |
| | Gourd | Cucumber, peeled, raw |
| | Pumpkin | Pumpkin, raw |
| | Spinach, frozen | Spinach, frozen, chopped or leaf, cooked, boiled, drained, without salt |
| | Mowa | Mustard greens, cooked, boiled, drained, without salt |
| | Pepper | Peppers, sweet, green, cooked, boiled, drained, without salt |
| | Taro | Taro, cooked, without salt |
| | Mubooora | Pumpkin leaves, cooked, boiled, drained, without salt |
| **Protein** | Beef, pork as main dish | Luncheon meat, pork, beef |
| | Birds | Goose, domesticated, meat and skin, cooked, roasted |
| | Chicken with skin | Chicken, broilers or fryers, back, meat only, cooked, fried |
| | Rabbit | Game meat, rabbit, domesticated, composite of cuts, cooked, stewed |
| | Mbeva | Mice, Rat |
| | Eggs | Egg, whole, cooked, hard-boiled |
| | Matemba | Fish, catfish, channel, wild, cooked, dry heat |
| | Sea snail | whelk, unspec, ckd,moist heat |
| | canned tuna | Fish, tuna, light, canned in water, without salt, drained solids |
| | Loucst | Locust |
| | Termites | Termites |

| | | |
|---|---|---|
| | Caterpillar | Caterpillar |
| **Grain** | White rice | Rice, white, long-grain, precooked or instant, enriched, prepared |
| | White bread | Bread, white, commercially prepared (includes soft bread crumbs) |
| | Potatoes | Potatoes, russet, flesh and skin, raw |
| | Dark bread | Bread, whole-wheat, commercially prepared |
| | Macaroni, cooked | Macaroni, whole-wheat, cooked |
| | sadza from maize | Corn, yellow |
| | sadza from sorghum | Sorghum |
| | Sadza from Millet | Millet, cooked |
| | Samp | Corn, sweet, white, canned, whole kernel, drained solids |
| | Sweet potato | Yam, cooked, boiled, drained, or baked, without salt |
| | Cassava | Cassava, raw |
| **Drinks** | Cascade | Cascade dairy fruit drink |
| | Soft drink, regular | Carbonated beverage, ginger ale |
| | Juice | Fruit juice, assorted raw |
| | Maheu | Alcoholic beverage, beer, regular, all |
| | Soft drink, light | Carbonated beverage, low calorie, other than cola or pepper, without caffeine |
| | Traditional beer | Alcoholic beverage, beer, regular, BUDWEISER |
| **Snacks** | Peanut butter | Peanut butter, smooth style, without salt |
| | Nuts | Peanuts, all types, raw |
| | Seeds | Seeds, pumpkin and squash seed kernels, roasted, without salt |
| | Cake, homemade | Cake, pound, commercially prepared, butter |
| | Cake, commercial | Cake, sponge, commercially prepared |
| | Biscuits | Biscuits - digestive |
| | Sweets | Candies, hard |
| **Fast Food** | Peri Peri Chicken | Nandos Peri Peri Chicken |
| | Fried Chicken | Chicken, broilers or fryers, leg, meat and skin, cooked, fried, batter |
| | Fries | BURGER KING, French Fries |
| | Hamburger | BURGER KING, Hamburger |
| | Pizza | PIZZA HUT 12" Pepperoni Pizza, Thick Crust |

**Table S2:** Food items in the Zimbabwean FFQ had USDA [45] food item equivalents for nutritional compositional references.
